# Parthenogenesis in weevils of the tribe Naupactini (Coleoptera, Curculionidae): a *Wolbachia*-density dependent trait?

**DOI:** 10.1101/2020.07.17.208447

**Authors:** Lucía da Cruz Cabral, Lucía Fernandez Goya, Romina V. Piccinali, Analía A. Lanteri, Viviana A. Confalonieri, Marcela S. Rodriguero

**Author notes:** Universidad Tecnológica Nacional, Facultad Regional Chubut (UTN-FRCh), Av. del Trabajo 1536, Puerto Madryn, Chubut, Argentina. Corresponding author (MSR).

## Abstract

The intracellular bacteria *Wolbachia pipientis* can manipulate host reproduction to enhance their vertical transmission. It has been reported an association between parthenogenesis and *Wolbachia* infection in weevils from the tribe Naupactini. A curing experiment suggested that a threshold density of *Wolbachia* is required for parthenogenesis to occur. The aim of this study was to analyze *Wolbachia* infection status in the bisexual species *Naupactus xanthographus* and *Naupactus dissimulator*.

*Wolbachia* infection was detected in both species from some geographic locations, not being fixed. In all positive cases, faint PCR bands were observed. Quantification through real time PCR confirmed that *Wolbachia* loads in bisexual species were significantly lower than in parthenogenetic ones; this strengthens the hypothesis of a threshold level. Strain typing showed that both species carry *w*Nau1, the most frequent in parthenogenetic Naupactini weevils. These infections seem to be recently acquired by horizontal transfer. *Wolbachia* was located throughout the whole body, which reinforce the idea of recent transmission. Moreover, we demonstrated that this strain carries the WO phage.

Finally, the analysis of eubacterial *16S rRNA* gene showed intense PCR bands for both bisexual species, suggesting –the presence of additional bacteria. Interspecific competition might explain why the parthenogenetic phenotype is not triggered.

## 1. Introduction

The obligate intracellular Gram-negative bacteria *Wolbachia pipientis* (Rickettsiales: Anaplasmataceae) is the most widespread endosymbiont in nature, infecting arthropods and filarial nematodes [1]. It is mainly transmitted vertically by females, but horizontal transfer is also extensive [2]. *Wolbachia* is able to manipulate host reproduction by inducing several disorders including cytoplasmic incompatibility, feminization of genetic males, embryonic and larval male killing and thelytokous parthenogenesis [3]. These reproductive alterations give a selective advantage to the bacterium, enhancing infection spread [1]. Nevertheless, *Wolbachia* can play plenty of roles in symbiotic associations, such as production of nutrients or resistance against pathogens [4]. It has been suggested that *Wolbachia* tissue localization may be important in relation to function [5]. Infection is not restricted to reproductive organs, different studies have detected that *Wolbachia* can colonize diverse somatic tissues such as muscle, digestive tract, brain, fat body and the hemolymph [5-10].

It has been stated that *Wolbachia*-induced host phenotypes are deeply influenced by bacterial titers [1,5,11-14]. For instance, a two-step mechanism of parthenogenesis reported for parasitoid wasps (i.e. diploidization of the unfertilized egg followed by feminization) occurs only if *Wolbachia* exceeds a density threshold within eggs [15]. However, complete understanding of the association between phenotype and *Wolbachia* density remains unclear.

*Wolbachia*-induced parthenogenesis (WIP) is only confirmed in host taxa with haplo-diploid sex determination, although it is suspected in a small number of diplo-diploid species, as springtails and weevils [16-20]. The strong overrepresentation of haplo-diploid species with WIP could be due to an ascertainment bias and may not necessarily represent a biological pattern [21]. This could be related to the difficulties to formally demonstrate WIP in taxa with diplo-diploid sex determination systems because curing experiments lead to sterility rather than to restoration of sexuality as in haplo-diploid species [16,18-20,22]. Most probably, the mechanism to ensure parthenogenetic reproduction is quite different from those proposed for wasps and thrips, e.g. egg diploidization via gamete duplication.

Considering its particular properties, *Wolbachia* is being studied as a potential tool for control of insect pests and pathogens of insect-borne diseases [5]. Therefore, it is considered a novel alternative to combine with other existing strategies for Integrated Pest Management in an environmentally friendly way [23]. So, any knowledge on new strains, like those inducing parthenogenesis in diplo-diploid arthropods, may be welcome in order to provide new ideas to develop more effective control strategies.

The tribe Naupactini comprises more than 500 species of weevils distributed mostly in Central and South America. Several species within this group reproduce parthenogenetically like *Naupactus cervinus* [24], and many others were proposed as presumably parthenogenetic on the basis of female- biased sex ratios such as *Pantomorus postfasciatus* [25]. It has been reported an association between this reproductive mode and *Wolbachia* infection, with parthenogenetic species or populations being *Wolbachia*-infected, while unisexual species or population are not [17,26]. The molecular mechanism behind this striking pattern is still unknown. However, a recent curing experiment performed on *P. postfasciatus* suggested that a threshold density of *Wolbachia* is required for parthenogenesis to occur [18]. Considering this, we wondered if the bisexual species previously described as uninfected were actually free of this bacterial infection. It is possible that bisexual species harbor lower densities than those required for triggering parthenogenesis. Indeed, bacterial loads may be too small to be detected by conventional PCR [27-29] and may have been unnoticed in the former survey of *Wolbachia* infection in the tribe Naupactini [17].

The aim of the present study was to analyze the *Wolbachia* infection status in two bisexual species from the tribe Naupactini, to compare *Wolbachia* density of these weevils with parthenogenetic hosts carrying the same strain, and to investigate *Wolbachia* tissue localization in both parthenogenetic and sexually reproducing species. The two bisexual species were selected considering their abundance in nature and their economic importance. *Naupactus xanthographus*, known as “the fruit weevil”, causes severe damage on peach, nectarine, apple, berries, cherries, and other deciduous fruit trees, as well as alfalfa, potatoes, soybean and other plants of commercial relevance [30]. Moreover, *N. xanthographus* causes damage in grape vineyards in the most productive areas of Chile, Argentina and Brazil. It is a quarantine pest in Japan and the USA [31] and several measures have been established to intercept this weevil from grape exports from Chile to Peru [32]. On the other hand, *Naupactus dissimulator* causes damage on other important commercial crops like citrus species, yerba mate tea (*Ilex paraguariensis* Saint Hill) and peach, among others [30]. This information may be useful to increase knowledge on the dynamics of some components of the microbiota of pest weevils as well as the factors that modulate bacterial density as potential tools for Integrated Pest Management.

## 2. Materials and methods

### 2.1. *Wolbachia* survey in bisexual species

#### 2.1.1. Sampling of biological material

Both male and female adult specimens of *N. xanthographus* were collected in 14 locations from Argentina during the summer season of 2004-18, while *N. dissimulator* individuals were sampled in 8 locations from Argentina and Brazil during the same period (S1 Table). *Wolbachia* infection status was determined in 1-2 individuals per location.

Samples were obtained using a beating sheet (0.55 x 0.55 cm). Specimens were stored at −20°C for DNA extraction.

#### 2.1.2. *Wolbachia* detection and strain typing

Total genomic DNA was extracted from adult weevils using the DNeasy Blood & Tissue Kit (Qiagen, Germany), following manufacturer instructions.

In addition, DNA from *Naupactus cervinus* and *Naupactus dissimilis* (sister species of *N. dissimulator* and *N. xanthographus*, respectively), both of them parthenogenetic and naturally infected with *Wolbachia* [17] were used as positive controls. Both parthenogenetic (infected) and bisexual (uninfected) populations of *P. postfasciatus* [26] were also included in this study. Distilled water was used as negative control.

Bacterial presence in weevils was assessed with eubacterial specific primers for the *16S rRNA* gene [33]. *Wolbachia* infection was diagnosed through different genomic regions, using specific primers of the cytochrome C oxidase subunit I (*coxA*), aspartyl/glutamyl-tRNA amidotransferase subunit B (*gatB*), and *Wolbachia* surface protein (*wsp*) [34-35]. Finally, primers S1718 and A2442 specific for the insect mitochondrial cytochrome C oxidase subunit I (*COI*) gene [36] were used to check the quality of the DNA extraction. Amplifications were carried out in a 15 μL final volume reaction containing 100 ng of genomic DNA used as template, 0.5 μM of each primer (Thermo Fisher Scientific, USA), 0.1 mM of each dNTP (GenBiotech, Argentina), 25 mM MgCl_2_ (Thermo Fisher Scientific, USA), 1 unit of Taq polymerase (Thermo Fisher Scientific, USA) and 1X buffer (Thermo Fisher Scientific, USA). The reactions were performed on an Applied Biosystems Veriti thermal cycler under the conditions described in [37] for the *COI* gene, [34] for the *coxA* and *gatB* genes, and [35] for the *wsp* gene. In the case of the *16S rRNA* gene, the thermal conditions were those described in [33], but using 50° C as annealing temperature. PCR products were run on a 1% agarose gel with TAE buffer and visualized using GelRed^®^ staining (GenBiotech S.R.L.). All experiments were repeated at least twice.

*Wolbachia* strains from *N. xanthographus* and *N. dissimulator* were characterized through amplification and sequencing of the *fbpA* gene, which is the most rapidly evolving of the five *Wolbachia* MLST genes, and then the most sensitive to detect the maximum diversity of *Wolbachia* strains [34,38-39]. Primers and thermal profiling were obtained from [34].

In addition, the *Wolbachia* infecting temperate phage was surveyed by sequencing the *orf7* locus from the WO phage. Primers and conditions described in [40] were used and reactions were performed in both bisexual species and in a parthenogenetic population of *P. postfasciatus.* In both cases PCR amplifications were carried out as formerly described, but T-Holmes Taq polymerase kit (Inbio Highway, Argentina) was used because of its higher sensitivity, which allows detection of DNA at very low levels. PCR products were enzymatically purified using Exonuclease I (ExoI) and Thermosensitive Alkaline Phosphatase (FastAP) (Thermo Fisher Scientific, USA). Sequencing in both directions were performed in a 3130-XL Automatic Sequencer (Applied Biosystems). Sequences of the *fbpA* gene were compared with the *Wolbachia* MLST website. BLASTN sequence analyses were conducted to identify the *orf7* sequences.

### 2.2. *Wolbachia* quantification in bisexual species

#### 2.2.1. Sampling of biological material

In order to measure *Wolbachia* levels in sexually reproducing Naupactini species and to compare these loads with that of a parthenogenetic control (*P. postfasciatus*), samples were collected during the summer season 2018-19. This parthenogenetic control was selected considering that the three species share the same *Wolbachia* strain (*w*Nau1). Two locations were selected for each species (four specimens each, two males and two females). *N. dissimulator*: Buenos Aires City (hereafter, CABA) and Paulino Island; *N. xanthographus*: CABA and Pereyra Iraola Park; *P. postfasciatus*: CABA and Colonia (Uruguay). Specimens were managed as previously described (section 2.1.1).

#### 2.2.2. Real Time Quantitative PCR

For relative quantification, *gatB* was used as target gene, while weevil’s *ITS1* was selected as endogenous control gene. A nested PCR was optimized in order to increase sensitivity and specificity of the method. For both genes, an end-point PCR was performed as a first step using 100 ng of total genomic DNA as template, followed by qPCR of the product.

End-point PCR was performed using the primers and thermal cycling conditions for *gatB* gene previously described (section 2.1.2). For the *ITS1* gene, the primers and conditions used were those described in [41].

Primers for *gatB* and *ITS1* genes suitable for qPCR analyses were designed using Primer3Plus software [42]. Primer sequences for *gatB* were based on the sequence of *w*Nau1 strain described in [17], whereas primer sequences for *ITS1* were designed using conserved regions of *ITS1* sequences from the three weevil species retrieved from GenBank (NCBI, NIH) (Table 1).

**Table 1.**
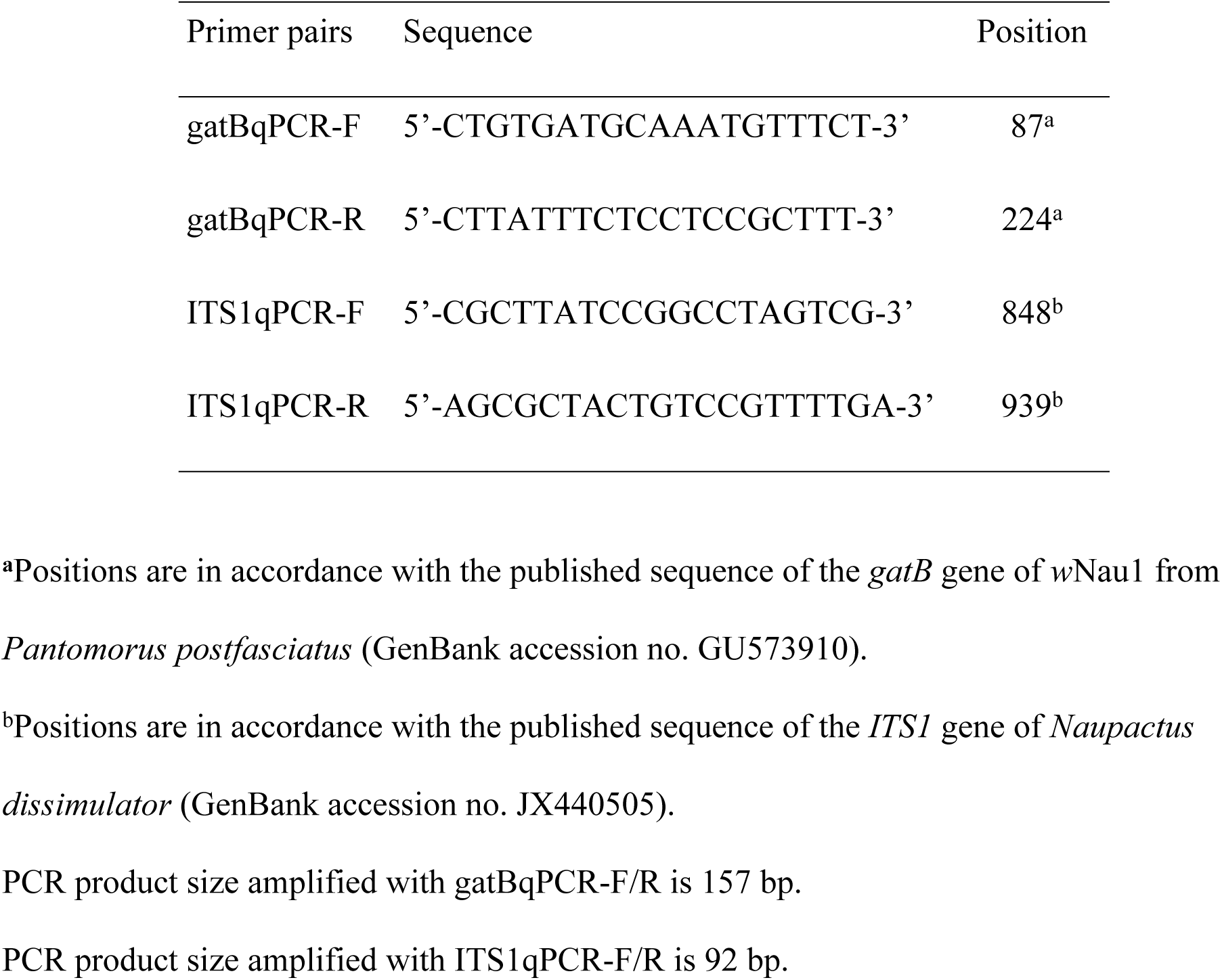
Oligonucleotide sequences of primers designed in this study for qPCR.

The qPCR reactions were conducted in a Step One Plus Real-Time PCR System (Thermo Fisher Scientific, USA) using the SYBR Green methodology. Reactions were prepared in 20 μL total volume mixtures, consisted of 10 μL SYBR™ Select Master Mix (Thermo Fisher Scientific, USA), 200 nM of each primer (Macrogen, Korea) and DNA template (1 μL of ITS1 or *gatB* PCR product 500-fold diluted). All qPCRs were run in triplicate and each run also included three replicates of a negative control with no added DNA template. The thermal cycling conditions for both genes were 95° C for 2 min followed by 40 cycles of 95° C for 3 s and 60° C for 30 s. After that, melting curve analyses of the PCR products were performed. Standard curves were constructed using a qPCR amplicon obtained for each species serially 10-fold diluted. The qPCR amplification efficiency was calculated from the formula E = (10^−1/S^), being S the slope of the linear fit in the standard curve [43]. The three species showed similar and adequately high efficiencies for both amplicons.

Relative *Wolbachia* levels were analyzed by the comparative Cq method [44], which standardizes target genes against an endogenous host gene and adjusts for differences in PCR efficiency between the amplicons, using the formula:

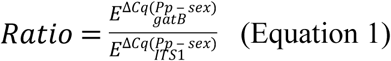

E= mean efficiency for each gene

Pp= mean Cq obtained for all *P. postfasciatus* individuals

sex= mean Cq obtained from the three replicates for each individual with sexual reproduction

#### 2.2.3. Data analyses

Differences in *Wolbachia* relative quantification were analyzed through general linear mixed models with the library lme [45], using RStudio v. 1.2.5033 [46] and R software environment v. 3.0.1 [47]. Normality of the data set was evaluated using the Shapiro-Wilk test, while homoscedasticity was evaluated graphically. In case of non-homoscedasticity, variance was modeled using varIdent. Both the Akaike Information Criterion (AIC) and Bayesian Information Criterion (BIC) were applied to select the best-fitting model. Analyses were carried out in two separate groups, including: (i) the whole dataset to evaluate the effect of the reproductive mode (n=24); (ii) *N. xanthographus* + *N. dissimulator* data to test the effect of the species, sex and location (n=16).

For (i), a model with the ratio obtained from Equation 1 as response variable, and reproductive mode as explanatory variable with fixed effects was applied. Geographic location was included as a random effect variable and was used to model variance.

For (ii), significance of explanatory variables (species, sex and location) was tested by dropping explanatory variables and their interactions from the models. Models considering the interactions among the explanatory variables did not fulfill the assumptions of normality and homoscedasticity, even after variance modeling. Sex was not considered in the final model because it has no significance (p>0.05) and both AIC and BIC values obtained were higher (AIC_sp+loc+sex_ = - 79.607 > AIC_sp+loc_ = −100.133; BIC_sp+loc+sex_ = −76.884 > BIC_sp+loc_ = −96.950). An additive model with the ratio obtained from Equation 1 as response variable, and species and geographic location as explanatory variables with fixed effects was selected. Variance was modelled by geographic location. PCR plate was used as explanatory variable with random effects.

All charts were performed with the R software package v. 3.0.1 [47], using RStudio v. 1.2.5033 [46].

### 2.3. *Wolbachia* tissue localization

#### 2.3.1. Biological material and dissection

Adults from the three species studied herein were sampled in CABA (5 *N. dissimulator* females, 5 *N. dissimulator* males, 3 *N. xanthographus* females and 3 parthenogenetic *P. postfasciatus*). They were conserved at −20° C after collection and then dissected with a scalpel using a stereo-microscope (100×). Each body was separated in four: head (H), reproductive tissue (R), digestive tissue (D) and rest of the body (B). Each sample was conserved in absolute ethanol at −20° C.

#### 2.3.2. DNA extraction and PCR

Total genomic DNA was extracted using the REDExtract-N-Amp™ Tissue PCR Kit (Sigma- Aldrich, USA), which yields high quantities of DNA. Samples were analyzed by end-point PCR for the *gatB* and the *16S rRNA* genes, as described in section 2.1.2. In addition, samples that resulted negative in agarose gel for the *gatB* gene were re-analyzed using qPCR for this gene, as explained in section 2.3.2. All experiments were repeated at least twice.

## 3. Results

### 3.1. *Wolbachia* survey in bisexual species

*Wolbachia* infection was detected in 8 out of 14 geographic locations investigated for *N. xanthographus* (Fig 1) for all the genes assayed. In addition, it was found in individuals of *N. dissimulator* from most of the locations surveyed (7 out of 8) (Fig 1).

**Fig 1.**
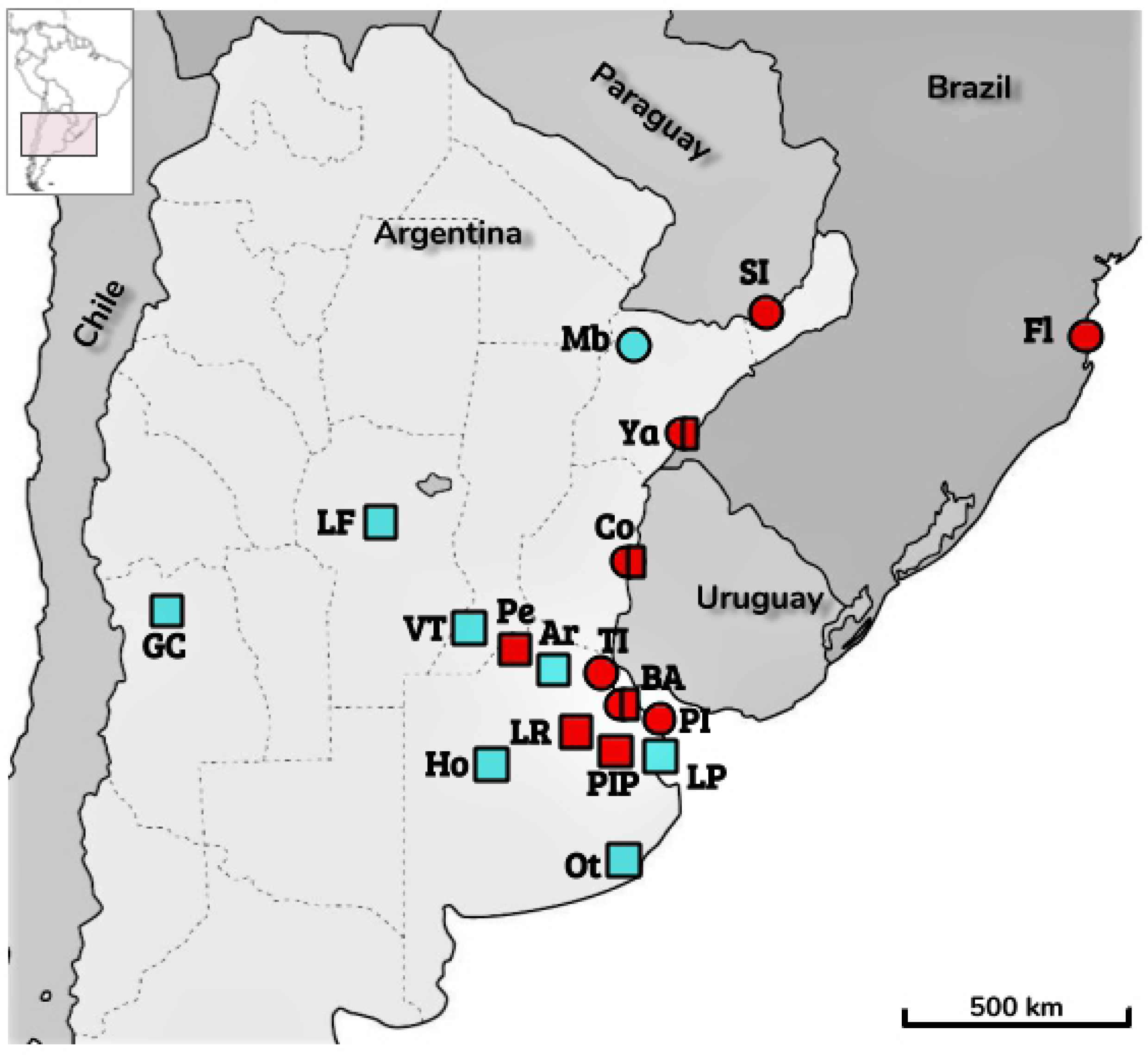
Distribution of sampling sites of *Naupactus dissimulator* (circles) and *Naupactus xanthographus* (squares). Colors indicate *Wolbachia* infection status at each sampling point (red, infected; blue, uninfected). For interpretation of the references to geographic locations in this figure, the reader is referred to the S1 Table.

In all cases, the bands observed in agarose gel were faint for both species, while parthenogenetic species used as positive controls showed intense bands for *Wolbachia* genes. Bisexual individuals of *P. postfasciatus* showed no *Wolbachia* DNA amplification (Fig 2A). On the other hand, the analysis of *16S rRNA* gene revealed the presence of intense bands for *N. xanthographus* and *N. dissimulator* (Fig 2B).

**Fig 2.**
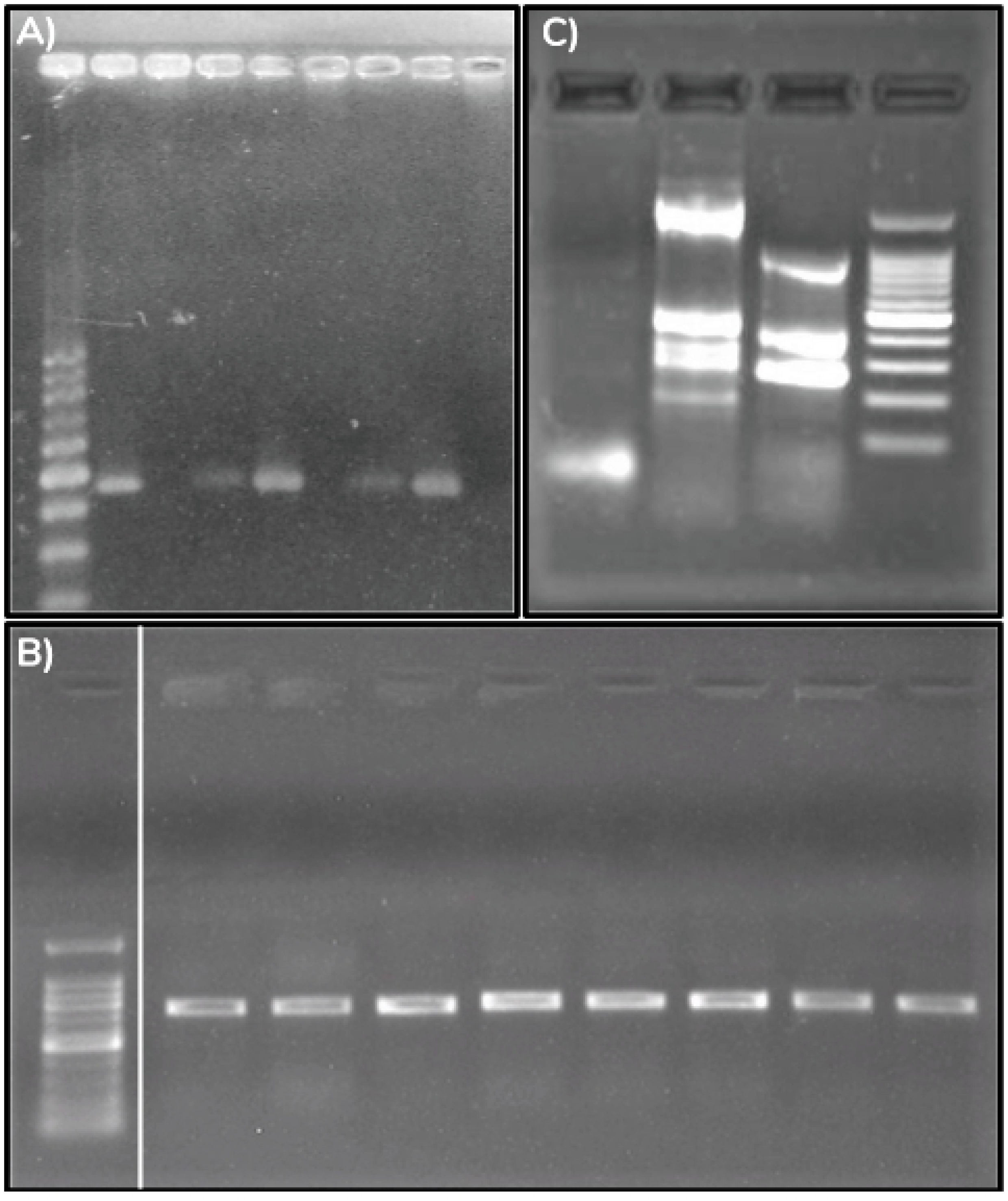
End-point PCR results in agarose gel. A) *Wolbachia gatB* gene; B) eubacterial *16S rRNA* gene; C) *orf7* locus from the *Wolbachia* WO phage. White line indicates a splicing in the gel. Lanes A and B: 1) DNA size marker; 2) *Naupactus cervinus* (parthenogenetic); 3) *Naupactus dissimulator* from Mburucuyá (bisexual, not infected); 4) *Naupactus dissimulator* from CABA (bisexual, infected); 5) *Naupactus dissimilis* (parthenogenetic); 6) *Naupactus xanthographus* from La Falda (bisexual, not infected); 7) *Naupactus xanthographus* from Colón (bisexual, infected); 8) *Pantomorus posfasciatus* (parthenogenetic); 9) *Pantomorus posfasciatus* (bisexual, not infected). Lanes C: 1) *Pantomorus posfasciatus* (parthenogenetic); 2) *Naupactus dissimulator* (bisexual, infected); 3) *Naupactus xanthographus* (bisexual, infected); 4) DNA size marker.

Sequencing of *fbpA* gene showed that both bisexual species have the 181 allele, which belongs to *w*Nau1 strain. The same strain was detected in most parthenogenetic populations of *P. postfasciatus* [26].

WO phage presence was revealed in the three species. While in parthenogenetic *P. postfasciatus* a single bright band in the gel was observed, both *N. dissimulator* and *N. xanthographus* showed multiple bands (Fig 2C). Sequencing of the *orf7* locus from *P. postfasciatus* confirmed identity with the WO phage (99.71% nucleotide identity with the WO capsid protein gene encoded by a *Wolbachia* strain from a butterfly native to India, GenBank accession no. FJ392499.1, E-value = 6e-174). The sequence obtained in the present study is available at GenBank (accession number MT526906).

### 3.2. *Wolbachia* quantification in bisexual species

The relative quantification obtained for *Wolbachia* in bisexual species was significantly lower than in parthenogenetic *P. postfasciatus* (Fig 3A).

**Fig 3.**
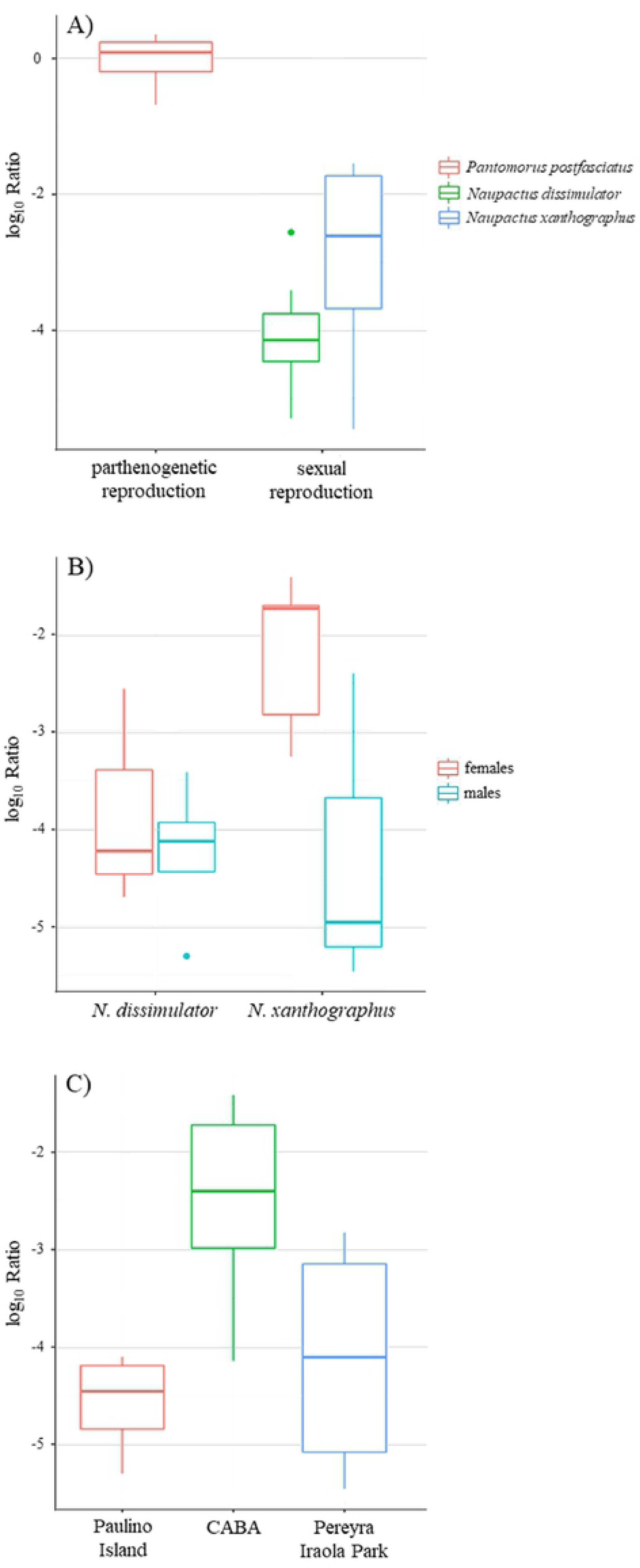
Relative *Wolbachia* levels (log_10_ ratio) for weevils with different reproductive modes: *Naupactus dissimulator* and *Naupactus xanthographus* (bisexual); and *Pantomorus postfasciatus* (parthenogenetic). Thick horizontal lines indicate median values, boxes the interquartile range (IQR), whiskers 1.5 times the IQR, and dots show outlier values. *Wolbachia* loads were compared by reproductive mode for the three species (A); sex split by bisexual species (B); and geographic location of bisexual species (C).

Thus, reproductive mode has a significant effect on *Wolbachia* levels (p=0.0331; estimated mean ratio for a bisexual population=1.73×10^−4^; DF=3).

The fold change due to reproductive mode was at least twice. Additionally, an elevated inter-individual variation in bacterial densities within bisexual species was detected.

When comparing within bisexual species, no significant effects were detected in *Wolbachia* loads related with species, sex or geographic location (p>0.05) (Fig 3B-C).

### 3.3. *Wolbachia* tissue localization in bisexual species

*Wolbachia* was detected throughout the different tissues analyzed in all the species and sexes surveyed. These results are summarized in Table 2.

**Table 2.** *Wolbachia* presence in different tissues in Naupactini species (*Naupactus dissimulator* and *Naupactus xanthographus* with sexual reproduction; and *Pantomorus postfasciatus* with parthenogenetic reproduction) expressed as proportion of positive samples/total of individuals analyzed. n: number of individuals; H: head; D: digestive tube; R: reproductive tissue; B: rest of the body.

|  | n | H | D | R | B |
| --- | --- | --- | --- | --- | --- |
| *N. dissimulator* female | 5 | 1 | 1 | 1 | 1 |
| *N. dissimulator* male | 5 | 0.6 | 0.6 | 0.6 | 0.6 |
| *N. xanthographus* female | 3 | 1 | 0.7 | 1 | 1 |
| *P. postfasciatus* female | 3 | 1 | 1 | 1 | 1 |

Even though infection was not detected in some tissues from a few samples, there were no individuals presenting the same pattern, indicating a random distribution of negative results, and then inferred as false negatives. Thus, the infection seems to be spread in the whole body of individuals with both reproductive modes studied. In addition, the analyses of the *16S rRNA* gene suggested the presence of other bacteria in the same samples considering that multiple bands were observed in the agarose gel (similar to the *orf7* locus).

## 4. Discussion

*Wolbachia* infection was detected in bisexual species from the tribe Naupactini at very low levels. Both *N. dissimulator* and *N. xanthographus* host *w*Nau1 strain. This is not surprising, since it is the most widespread within the tribe Naupactini, including *Pantomorus postfasciatus, Naupactus minor*, among other species [17,26]. Then, *w*Nau1 has an elevated incidence rate and is efficiently transmitted within this tribe. These infections appear to be not fixed, since they were recorded only in some geographic locations for the two bisexual species, revealing a much more complex host- symbiont relationship than previously thought [17]. Similar low *Wolbachia* infection frequencies and low bacterial densities were described for populations of the bark beetle *Pityogenes chalcographus* (Curculionidae, Scolytinae) [27] and *Drosophila melanogaster* [48-49].

In *P. postfasciatus*, a species with both and parthenogenetic bisexual populations, *Wolbachia* infection is not fixed either, and the strain *w*Nau1 is spatially scattered (see Figure 1 in [26]). Contrary, in *N. xanthographus* and *N. dissimulator Wolbachia* infection appears to be uniformly widespread throughout the geographic range of both species. The pattern found stimulates to deepen the study on *Wolbachia* dynamics in these species.

Additionally, we report for the first time the presence of the temperate phage WO in Naupactini weevils: *N. xanthographus, N. dissimulator* and *P. postfasciatus* yielded a positive diagnosis. Although the three of them share the same *Wolbachia* strain, their phages seems to be non-identical, as the agarose gel revealed a different band pattern between parthenogenetic and bisexual ones. Bordenstein et al. suggested that low *Wolbachia* densities might be caused by high densities of its associated bacteriophage [50]. Considering our results, this hypothesis deserves to be tested. Further studies will be conducted to deepen knowledge about phage WO in Naupactini.

*Naupactus xanthographus* is currently widespread in Argentina, whereas *N. dissimulator* is restricted to the gallery forests of Paraná and Uruguay rivers, down to the banks of La Plata River [30]. In this area, *N. dissimulator* coexists with its probable sister species, *N. cervinus* [37]. However, they do not share the same endosymbiont strain; *N. dissimulator* is infected with *w*Nau1 strain, while *N. cervinus* carries *w*Nau5. Something similar occurs with the sister species pair *N. xanthographus*- *N. dissimilis* (carrying *w*Nau1 and *w*Nau7, respectively [17]). This suggests that both *w*Nau1 and *w*Nau5 (or *w*Nau1 and *w*Nau7) were independently acquired. Coexistence of infected and uninfected populations also point to incipient, still-evolving processes, opposite to what was observed for older infections like that of *N. cervinus*, in which *w*Nau5 is fixed [51]. Furthermore, these infections seem to be recent since *w*Nau1 is shared for many distantly related hosts [17]. Multiple mechanisms of *Wolbachia* horizontal transmission have been proposed, including predators, parasitoids, hemolymph transfer, cohabitation, and foraging on the same host plants [52-55]. So far, in the case of weevils, mainly indirect evidence was provided of such transmissions [56-59].

Being primarily vertically transmitted, it is expected that the localization of *Wolbachia* would be restricted to the reproductive tissues. However, according to our results, these bacteria were not circumscribed to a specific tissue; instead, they are distributed throughout the whole body both in the bisexual and the parthenogenetic species studied. These results reinforce the idea of recent horizontal transfer and are in agreement with previous observations in ants [60] and mosquitoes [61].

As it was mentioned before, the same *Wolbachia* strain was detected in both bisexual and unisexual species under study, but at significant lower densities in the former ones. No difference was found in bacterial titers between *N. dissimulator* and *N. xanthographus*, and regarding sex or geographic location, although they presented higher variability in *Wolbachia* loads among individuals than parthenogenetic *P. postfasciatus*. These results strengthen the hypothesis of a threshold level for *w*Nau1 strain, i.e. a minimum bacterial load necessary to induce the parthenogenetic phenotype. Several studies have pointed out the importance of a quantitative measure of *Wolbachia* to correlate the effects on host manipulation [5,11,15,62-64]. However, there are few reports specifically evaluating the relationship between *Wolbachia* titers and the parthenogenetic phenotype, and all of them are referred to Hymenoptera [65-68]. To the best of our knowledge, our contribution would be the first report for Coleoptera.

The pattern observed for the bisexual species would suggest that some populations are evolving towards endosymbiont loss. Alternatively, they could constitute an example of a persistent *Wolbachia* infection at low levels and frequencies, as in the bark beetle *P. chalcographus* [27]. Several studies have stated that microbial symbionts in eukaryotes are not transient passengers randomly acquired from the environment [3,69-72]. Then, a question that remains is which function has *Wolbachia* in bisexual species. It is likely that a yet unidentified beneficial fitness effect conserves this infection under certain environmental conditions, even at these low levels [27].

Another interesting topic is why the parthenogenetic phenotype is not triggered in these species. A possible explanation for the association between *Wolbachia* and parthenogenesis is the lesser ability of parthenogenetic weevils to rid themselves of *Wolbachia* infections once these happen [17,19]. Actually, it appears that no species is able to dispose of *Wolbachia*. Instead, as it was demonstrated in the present work, bisexual species are able to maintain the infection at low densities. Then, they must have a mechanism to deal with such infection and consequently to impede parthenogenesis induction. Which could be this underlying mechanism?

First, *Wolbachia* levels might be modulated by proteins produced by the host. So far, several loci, either prokaryotic or eukaryotic, are known to play a role in the *in vivo* modulation of *Wolbachia* titer. This was demonstrated with experiments using hybrid hosts of the tsetse fly that strongly suggest that infected animals are actively controlling *Wolbachia* population dynamics [73]. In addition, another key inquiry is what environmental factors influence endosymbiont density. Among them, aspects related to host and endosymbiont metabolic and signaling pathways involved in nutrient sensing might be affecting *Wolbachia* levels [74]. These unexplored areas will be the origin of new research lines.

Another possible explanation is the presence of some other bacteria competing with *Wolbachia*, as suggested by the results obtained for the *16S rRNA* gene, i.e. bright bands, hint of many other bacteria besides *Wolbachia* (Fig 2). Symbioses are determined by highly dynamic interactions, both between the host and its symbionts, and among the different members of the symbiotic community [75]. It has been reported that conflict or incompatibility among microorganisms within arthropods, e.g. through competition for resources or space within the shared host, can shape their microbiome composition and could be a potential barrier to transmission of heritable symbionts [76]. Alternatively, particular taxa could provoke a host immune response, which in turn might affect the complete microbiota. There are few reports about interactions between *Wolbachia* and other bacteria, except for several binary interactions with other highly abundant symbionts. For instance, Goto et al. showed that male-killing *Spiroplasma* negatively affect *Wolbachia* titers in *D. melanogaster* [77], while Hughes et al. reported a mutual competitive exclusion between *Wolbachia* and *Asaia* in the reproductive organs of *Anopheles* mosquitoes [76]. Likewise, some still unidentified components of the microbiota in the bisexual species herein studied may out-compete *Wolbachia* either by a more efficient use of resources or by the production of some metabolite capable of regulating their levels.

It is possible that toxic effects, if any, of these unknown bacteria may provide clues to novel microbicidal mechanisms that may out-perform antibiotics. Under this hypothesis, these still unidentified components of the microbiota of species like *N. dissimulator* and *N. xanthographus*, should be absent in other species, such as *P. postfasciatus* and *N. cervinus*, for example, which display parthenogenesis at least in part of their ranges. This lack of specific bacteria could allow *Wolbachia* to proliferate and thereby surpass the threshold density in the latter species. Altogether, our findings add further support to the hypothesis of WIP in Naupactini weevils.

*Wolbachia*-microbiota interactions may be complex and dependent on both host and microbial composition. Future studies of high-throughput sequencing of the *16S rRNA* gene will be the starting point to test this hypothesis in both bisexual and unisexual species from Naupactini and to explore the microbiota composition of South American weevils.

## 5. Acknowledgements

Thanks are due to Dr Noelia Guzmán for *Naupactus xanthographus* DNA, to Dr Lucía Babino for statistical advice, and to Dr Agustin Elias-Costa for his valuable suggestion for designing Fig 2.

## 7. Supporting information

**S1 Table**: **Geographic distribution of *Naupactus dissimulator* (Nd) and *Naupactus xanthographus* (Nx) sampling points**. Acronyms of locations shown in Fig 1, coordinates and presence of *Wolbachia* infection in weevils sampled are also presented for each site.

